# Subchondral bone and synovial fluid metabolomic profiles are altered in injured and contralateral limbs 7 days after non-invasive joint injury in skeletally-mature C57BL/6 mice

**DOI:** 10.1101/2022.04.16.488558

**Authors:** Brady Hislop, Connor Devine, Ronald K. June, Chelsea M. Heveran

## Abstract

**Objective:** Post-traumatic osteoarthritis (PTOA) is a common long-term outcome following ACL injury. However, early changes to bone and synovial fluid after ACL injury are not sufficiently understood. The objectives of this study were to (1) evaluate whether acute bone loss one week after ACL injury is accompanied by altered subchondral bone plate modulus, (2) determine if bone changes are localized to the injured limb or extend to the contralateral-to-injured limb compared with sham-loaded controls, and (3) identify shifts in synovial fluid metabolism unique to injured limbs.

**Design:** Female C57Bl\6N mice (19 weeks at injury) were subjected to either a single tibial compression overload to simulate ACL injury (n=8) or a small pre-load (n=8). Mice were euthanized 7 days after injury, and synovial fluid was immediately harvested for metabolomic profiling. Bone microarchitecture, bone formation, and subchondral bone modulus at the proximal tibia were studied using microCT, histomorphometry, and nanoindentation, respectively. Osteoclast number density was assessed at the distal femur. For each bone measure a mixed model ANOVA was generated to determine the effects of injury and loaded side.

**Results:** Epiphyseal and subchondral bone microarchitecture decreased while subchondral bone tissue modulus was unchanged after ACL injuries. Bone resorption increased but bone formation was not changed. Loss of bone microarchitecture also occurred for the contralateral-to-injured limb, demonstrating that the early response to ACL injury extended beyond the injured joint. While the metabolomic profiles of the injured and contralateral-to-injured limbs had many similarities, there were also distinct metabolic shifts present in only the injured limbs. The most prominent of the pathways was cysteine and methionine metabolism, which is associated with osteoclast activity.

**Conclusion:** These results add to the understanding of early bone changes following ACL injury. Confirming prior reports, we observe a decline in epiphyseal and subchondral bone microarchitecture. We add the finding that subchondral bone modulus remains unchanged at one week after ACL injury. A potential biomarker of this initial bone catabolic response may be synovial fluid cysteine and methionine metabolism, which was only dysregulated in injured knees. Our results implicate a rapidly changing biological and mechanical environment within both the injured and contralateral joints that has the potential for influencing the progression to PTOA.

## 1.0 Introduction

Post-traumatic osteoarthritis (PTOA) affects 5.6 million people in the US each year^1^. Patients who suffer ACL tears develop PTOA at an alarming rate. Regardless of ACL reconstruction status, 50% of patients will develop PTOA within 20 years of joint injury^2^. Early interventions that mitigate the first deleterious changes to joint tissues may relieve some of the immense physical and financial burdens of PTOA^3–5^. However, to design and implement early interventions, additional data are needed to improve the understanding of how the loading and biological environments of the injured knee change early after joint injury.

It is well-established that subchondral and epiphyseal bone undergo transient thinning after joint injury that later reverses and progresses towards sclerosis^6–10^. However, because changes to subchondral bone modulus are not known, the impacts of early bone changes to the bone mechanical environment are not sufficiently understood. The rapid decline of subchondral bone and epiphyseal microarchitecture may directly impact the bone mechanical environment. If subchondral and epiphyseal bone microarchitecture decreases, µstrain will increase in subchondral bone with potential impacts to bone mechanotransduction^11,12^. The changes would be expected to be further exacerbated by changes to tissue-scale subchondral bone modulus, since the stiffness of bone tissue influences the strain experienced by osteocytes, a bone celltype recently implicated in the development of PTOA^13,14^. However, it is unknown whether subchondral bone modulus changes after joint injury. Thus, it is critical to quantify subchondral bone modulus changes after joint injury to better understand the effects of early bone changes on the bone mechanical environment.

The impacts of joint injury may extend to the contralateral-to-injured limb. Patients who undergo an initial knee replacement have a 37.2% chance of needing a contralateral limb replacement within 10 years^15^. In addition, Christiansen and coauthors found decreased trabecular microarchitecture in the L5 vertebra 10 days after non-invasive joint injury in mice^16^. Therefore, decreased bone microarchitecture may also acutely occur in the contralateral-to-injured knee. This bone loss may be the consequence of altered loading^17^, a systemic inflammatory response^18,19^, and/or altered remodeling bone cell populations^13,20,21^. However, rodent models of PTOA commonly only compare early outcomes after joint injury between injured and contralateral-to-injured limbs^8,10,22,23^. Comparisons between injured and sham-injured limbs have been limited to later time points when bone is sclerotic and PTOA is established^8,22,23^. Therefore, important gaps remain in our understanding of early bone changes in the injured and contralateral-to-injured limbs.

Synovial fluid interacts with multiple joint tissues including bone and cartilage. Therefore, shifts in synovial fluid metabolism marked by changes in metabolites could point to specific biological changes that rapidly occur after ACL injury in the injured joint. These potential biomarkers may have high utility in the design of early interventions to mitigate the progression to PTOA. Metabolomics measures metabolites (i.e., small molecules chemically changed in metabolism), providing a snapshot of the metabolic state of a fluid, tissue, organ, or system^24^. Since bone is highly cellular and vascularized compared with cartilage, assessing metabolic shifts in synovial fluid is expected to produce data that can help contextualize early bone changes after joint injury. Recent studies find distinct metabolic profiles between injured and sham-injured synovial fluid 7d after joint injury in mouse models^25,26^. However, the relevance of shifts in synovial fluid metabolism in injured versus sham-injured, and contralateral-to-injured soon after joint injury on bone outcomes remains uncertain.

In this study, we address several critical gaps about the early response of the injured joint to ACL injury. We determine the response of subchondral bone modulus and microarchitecture 7d after joint injury in a non-invasive murine model. We also compare early bone microarchitecture changes and synovial fluid metabolic shifts between the injured, contralateral-to-injured, and sham-injured limbs.

## 2.0 Materials and Methods

### 2.1 Mouse model of joint injury

All animal procedures were approved by the Montana State University Institutional Animal Care and Use Committee. C57Bl\6N mice were acquired from Charles River laboratories. Mice were allowed to acclimate for two weeks prior to injury (19-weeks old at injury). Each mouse was assigned to either the injury (overload-induced ACL tear) or control (sham-injury) groups. The injured or sham-injured joint was randomly assigned to a side.

On the day of injury, mice were anesthetized in an inhalation chamber (2-4% isoflurane, ∼30 seconds) and then nose cone (2% isoflurane) while one limb was loaded into a custom-built loading machine (**Figure 1A-B**). Sham-injured and injured limbs were subjected to a pre-load (1-2N, 0.5mm/s). Injured limbs were loaded to 12-15N at 130 mm/s, consistent with prior studies (**Figure 1C**)^7,8,28,29^.

**Figure 1.**
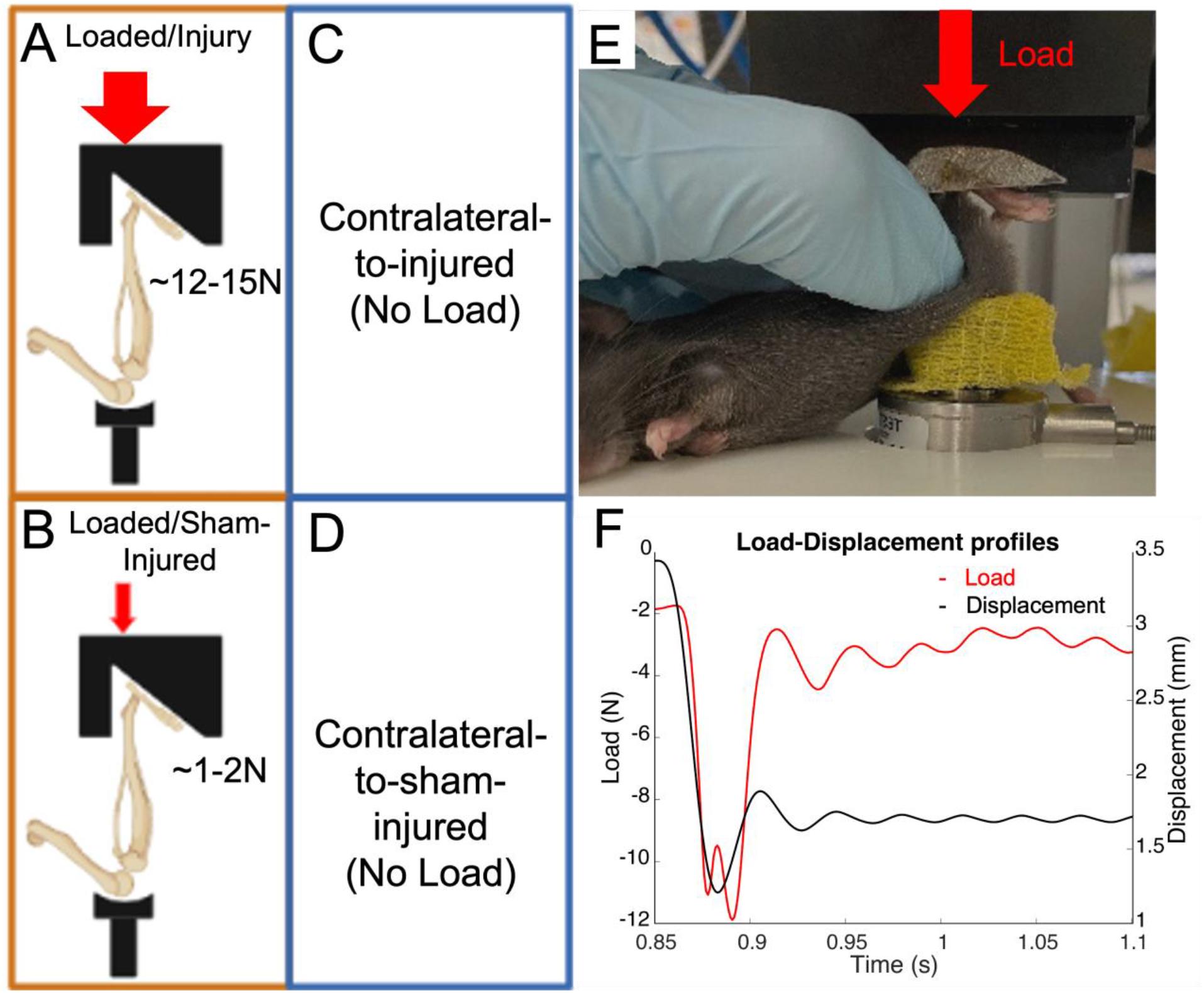
Mouse PTOA model. A, B) Schematic of mouse hind limb loaded into loading machine with load applied parallel to tibia, C,D) Contralateral limbs of both injured, and control mice were not loaded, E) Mouse loaded into loading machine during study, F) Load-displacement curves showing a change in displacement coinciding with a sinusoidal load response between 0.875- and 0.9-seconds characteristic of an anterior translation of the tibia leading to ACL rupture^8,28,29^. A,B) Created with Biorender.com

ACL tears were confirmed *in vivo* via the presence of the characteristic release in compressive load, and by performing pre- and post-injury joint laxity tests of anterior stability^7,8,28^. Five mice were excluded for failed ACL tears and two were excluded due to tibial fractures. Sixteen mice (n=8 injured, n=8 sham-injured) were studied for subchondral bone response and synovial fluid responses to joint injury.

Mice were group housed (≤5 mice per cage) and provided rodent diet (Purina LabDiet 5053) and water *ad libitum*. Calcein (20 mg/kg) and alizarin (30 mg/kg) fluorochrome labels were administered intraperitoneally 2-3 days before injury and 6 days post-injury, respectively. Mice were euthanized 7 days post-injury *via* cervical dislocation.

### 2.2 Analysis of trabecular and subchondral bone microarchitecture of the proximal tibia

Tibiae were cleaned of non-osseous tissue, submerged in 70% ethanol, and refrigerated (4°C). Proximal tibiae were evaluated for microarchitecture of the epiphysis and subchondral bone (μCT40, Scanco Medical AG, 10 μm isotropic voxel size, 70 kVP, 114 μA, 200 ms integration time). The scanned volume was a 1 mm (100 coronal slices) thick region at the center of the proximal tibial epiphysis. The first region of interest included the epiphysis (**Figure 2C**). Analyses included bone volume (BV, mm^3^), total volume (TV, mm^3^), bone volume fraction (BV/TV) (%), bone mineral density (BMD) (mgHA/cm^3^), trabecular number (Tb.N, 1/mm), trabecular spacing (Tb.Sp, mm), and trabecular thickness (Tb.Th, mm)^30^. The second region of interest included both the epiphysis and subchondral bone plate (**Figure 2C**). This region was assessed for BV, TV, BV/TV, tissue mineral density (TMD, mgHA/cm^3^), and BMD. A segmentation threshold of 500 mg/HA cm^3^ was used to segment bone from soft tissue prior to analysis.

**Figure 2.**
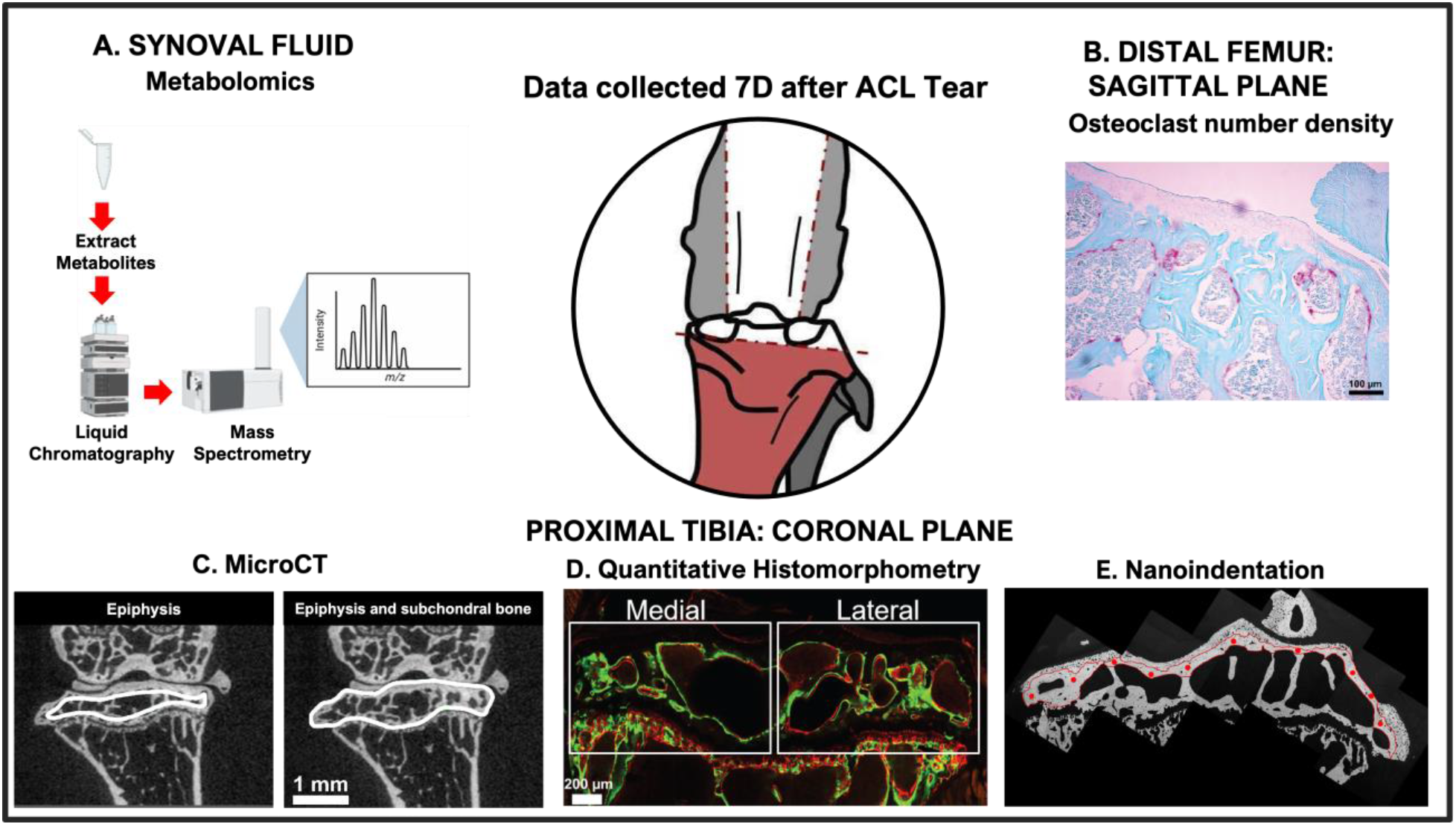
Schematic of experimental methods to characterize bone and synovial fluid responses to joint injury. A) Schematic of synovial fluid metabolomic extraction and analysis procedure from extraction to mass spectra. B) The osteoclastic activity at the distal femur was evaluated from TRAP-stained sections. C) Microarchitecture of the epiphysis and epiphysis and subchondral bone were evaluated from MicroCT. D) Dynamic bone remodeling for the epiphysis was measured from confocal laser scanning microscopy images from polished coronal sections. Medial and lateral regions of interest are delineated by the white boxes. E) Subchondral bone modulus was measured using nanoindentation on polished coronal sections. Indents were placed within the outlined region spanning the medial to lateral aspects of the subchondral bone plate. A) Created with Biorender.com

### 2.3 Evaluation of osteoclastic bone resorption at the femoral epiphysis

Femurs were placed in 10% neutral buffered formalin (NBF) for 18 hours, decalcified in Ethylene Diamine Tetraacetic Acid (EDTA), dehydrated in a graded series of ethanol concentrations (70-100%), embedded in paraffin, and microtomed to 6µm-thick sagittal sections. Sections were stained via standard protocols for tartrate resistant acid phosphatase (TRAP) and imaged using a brightfield microscope (Nikon, Eclipse E-800) (**Figure 2B**). Using ImageJ,^31^ an evaluator blinded to joint injury status traced the epiphysial trabecular bone surfaces enclosing the marrow cavities (4X images) and counted the multinucleated, TRAP-positive cells (*i*.*e*., dark pink) (10X images). We report the osteoclast number density normalized to marrow area and bone surface.

### 2.4 Analysis of dynamic bone remodeling of the epiphysis

Following microCT, tibiae were embedded in poly(methyl)methacrylate, sectioned along the coronal plane of the tibial plateau, and polished to a mirror finish as previously described (**Figure 2**)^32^. Calcein and alizarin labels were imaged at the proximal tibia using confocal laser scanning microscopy (CLSM, inverted Leica SP5). Samples were excited at 488 nm and 561 nm. Emission windows were set at 500 nm-550 nm (calcein) and 650 nm-750 nm (alizarin). The voxel size was 1.52µm by 1.52µm by 4.74µm. Z-stack was collected for each sample in medial and lateral regions. One in-focus slice from each z-stack was identified for analysis (Imaris 9.3.0, Bitplane). The original images were thresholded until the boundaries of the trabeculae were identifiable for measurement of bone perimeter. The threshold was then removed, and the contrast of the original image was set to a maximum for measurement of bone labels.

Bone perimeter and calcein and alizarin labels were traced on iPads by two independent evaluators blinded to joint injury status. If inter-evaluator agreement was less than 95%, sections were re-analyzed by both evaluators. Bone perimeter and labels were measured in epiphyseal regions of interest (**Figure 2D**). For each region of interest, bone surface, alizarin surface / bone surface, and calcein surface / bone surface were calculated. Each measure was calculated using a custom written MATLAB script (available at: https://github.com/hisl6802/Quantitative-Histomorphometry-Automated-Tracing-).

### 2.5 Evaluation of tissue-scale subchondral bone plate modulus

Nanoindentation was performed on subchondral bone from polished proximal tibia sections using a KLA Tencor iMicro (5 mN max load, 30s load and unload, 60s hold) and Berkovich tip (ν=0.07, E=1140 GPa). Indents were placed between the tidemark and the epiphysis spanning the medial to lateral subchondral bone plate (n=8 per sample) (**Figure 2E**). Indentation curves were analyzed for reduced modulus (E^r^) consistent with Vahidi et al^32^. We report the indentation modulus E^i^, which is corrected for the diamond tip properties (E^t^, ***υ***^t^), but avoids assuming a Poisson’s ratio for the tissue sample (***υ***^s^) (**Equations 1-2**)^33^.

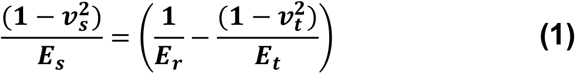

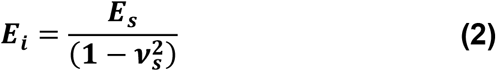

Indents were excluded in circumstances of poor surface contact, a non-smooth loading or unloading curve, or moduli indistinguishable from PMMA (≤5 GPa).

### 2.6 Profiling the biological response of joints to traumatic injury

Synovial fluid was extracted from the stifle joint immediately following euthanization consistent with our previously published protocol^26^. Briefly, an incision was made anteriorly into the synovium and synovial fluid was collected by adsorption using Melgisorb alginate dressing (Tendra). Metabolites were extracted as previously described^34,35^ and analyzed using Liquid Chromatography-Mass Spectrometry (LC-MS, Agilent 1290 UPLC, Agilent 6538 Q-TOF mass spectrometer) (**Figure 2A**). Spectra were submitted to XCMS^36^ for peak identification.

### 2.7 Data Analysis

#### 2.7.1 Evaluation of the effects of injury and loaded side on bone outcomes

Measures of bone microarchitecture, labeled bone surfaces, osteoclast number, and nanoindentation modulus were analyzed using mixed model ANOVA. Fixed effects included injury (injury vs sham-injury) and load side (loaded and sham-loaded versus contralateral). The individual mouse was a random effect. For nanoindentation modulus, which included multiple indents per sample, the relative distance from the medial edge was included as a covariate. For fluorochrome label assessments, location (e.g., medial) was a fixed effect and limb was a random effect nested within the individual mouse.

Transformations of the dependent variable were performed, if necessary, to satisfy ANOVA assumptions of residual normality and homoscedasticity. Critical alpha was set *a priori* at 0.05. Significant interactions were tested for simple effects while maintaining family-wise error at 0.05 using a Bonferroni correction. Spearman rank correlation analysis was performed to evaluate relationships between significant changes with respect to injury in bone microarchitecture versus peak load.

#### 2.7.2 Metabolomic analysis

ANOVA (analysis of variance), t-tests, principal component analysis (PCA), partial least squares-discriminant analysis (PLS-DA), and hierarchical clustering were used to compare metabolic profiles for synovial fluid from the injured, sham-injured, and contralateral-to-injured limbs. The metabolic data were then subjected to cluster validation and ensemble clustering to determine the metabolites which consistently clustered together (**Supplement**) (**Figure 3**). The 10 largest clusters identified through ensemble clustering were submitted to mummichog to identify the associated metabolic pathways^37^.

**Figure 3.**
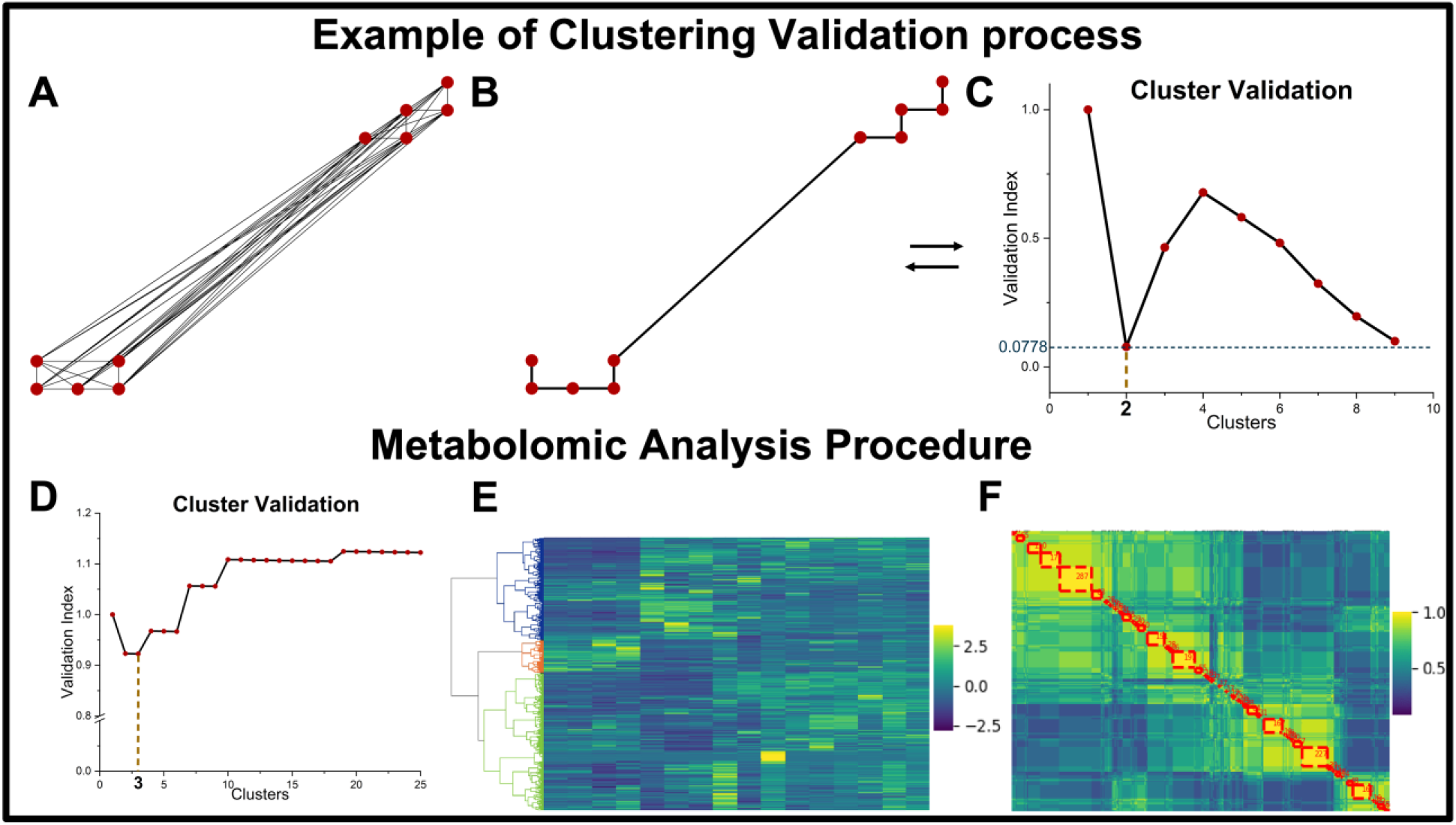
Example of minimum spanning tree (MST) based cluster validation. Metabolomic clustering analysis procedure. A) Ten 2D points (randomly generated using MATLAB) connected in a full dissimilarity tree; B) MST for the ten points; C) Cluster validation for B showing that 2 clusters (0.0778 validation index) is the optimum choice. D) Cluster validation output for analysis of Sham-injured v. Injured limb metabolic profiles showing a local minimum at 3 clusters; E) Graphical representation of one of thirteen hierarchical clustering outputs of the sham-injured v. injured metabolomic data, with the three clusters of interest colored (blue, orange, and green); F) Ensemble clustering output showing the clusters with greater than 40 metabolites in red dashed squares (yellow represents metabolites which clustered together in each of the thirteen clusterings).

## 3.0 Results

### 3.1 Epiphyseal and subchondral bone plate microarchitecture of the proximal tibia declines 7d after joint injury

MicroCT analysis of the tibial epiphysis revealed significant effects of injury (i.e., injury versus sham-injury) and loaded side (i.e., the injured and sham-injured limb versus the contralateral limbs) on several measures of bone microarchitecture. Epiphyseal BV/TV decreased with injury (−10.8%, p=0.002) and loaded side (−7.5%, p=0.023). Epiphyseal BMD also decreased with injury (−7.1%, p=0.012) and loaded side (−5.8%, p=0.040). Similarly, epiphyseal Tb.Th decreased with injury (−5.5%, p=0.028) and loaded side (−6.0%, p=0.017). For each of these measures, there were no significant interactions. TB.N and Tb.Sp were not affected by injury, loaded side, or the interaction of these factors. Because the peak load to cause the joint injury varied between mice, we tested whether peak load influenced measures of bone microarchitecture that had significant effects of injury. Spearman’s rank correlation analysis of peak load vs. BV/TV, BMD or Tb.Th found no significant relationships (p>0.05 for both).

A larger region of interest including subchondral bone and the epiphysis also showed significant decreases in BV/TV and BMD with injury (−5.8%, p=0.0016; −4.6%, p=0.012, respectively) and loaded side (−3.9%, p=0.030; −3.8%, p=0.037, respectively) (**Table 2**). The magnitude of bone loss in this region was less than for the epiphysis alone. There was a significant interaction between injury and loaded side on TMD (p=0.0415). However, post-hoc testing found no significant differences in TMD between injured and contralateral-to-injured (p=0.118) nor sham-injured and contralateral-to-sham (p=1). There was not a significant relationship between peak load and BV/TV or BMD.

### 3.2 Osteoclastic resorption at the distal femoral epiphysis increases but proximal epiphysis bone formation is unchanged 7d after joint injury

Injury significantly affected both osteoclast area number density and osteoclast number per bone surface at the distal femur epiphysis (+287.2%, p=0.017; +267.4%, p=0.016, respectively) (**Table 3**). These measures were not impacted by loaded side nor the interaction between loaded side and injury. Thus, there was a similar increase in osteoclast abundance for both injured and contralateral-to-injured limbs.

We performed histomorphometric analysis of the proximal tibial epiphysis to determine whether bone formation was affected by joint injury. A calcein label was administered 2-3 days before the injury. There was a significant effect of loaded side on calcein-to-bone-surface (−18.3%, p=0.005), indicating that bone resorption was increased for loaded (injured and sham-injured) limbs. Calcein labeling was not affected by injury, region of interest (e.g., medial), or the interaction of these factors. Alizarin was administered 6 days after the joint injury and 1 day before euthanasia. Alizarin-to-bone-surface was not affected by injury, loaded side, region of interest, or the interaction of these factors. Therefore, bone formation surface was not impacted by the joint injury.

### 3.3 Tissue-scale subchondral bone plate modulus is not changed 7d after injury

Nanoindentation tested whether subchondral bone plate modulus changes soon after joint injury. We first evaluated whether indent position (i.e., relative distance from the medial edge) influenced the indentation modulus (E^i^). Since a covariate describing location within the subchondral bone plate was not significant (p=0.057), we used a mixed model ANOVA without the covariate to test the effects of injury and loaded side on E^i^. There were no significant effects of injury (p=0.052), loaded side (p=0.338), nor an interaction between these factors (**Table 3**).

### 3.4 Distinct metabolic changes are evident in synovial fluid from injured limbs 7d after joint injury

We compared synovial fluid metabolic profiles from injured, sham-injured, and contralateral-to-injured limbs. A total of 3402 metabolites were detected across all synovial fluid samples, five metabolites were considered undetected and removed (zero median intensities for each group).

#### 3.4.1 Comparison of injured, sham-injured and contralateral-to-injured reveals dysregulated metabolites

One-way ANOVA detected significantly dysregulated metabolites. PCA, PLS-DA and hierarchical clustering detected only minimal group differences. However, some of these analyses (i.e., hierarchical clustering) are inherently subjective due to the sparsity of the data^38^. Therefore, we employed *ensemble clustering*^39^ to remove the subjectivity from metabolite clustering. Before cluster analyses, the median intensity of each metabolite in each group was determined. From this, clustering validation found that 9 clusters were optimum for these analyses. Ensemble clustering (**Supplementary Figure 1**) combined with the mummichog algorithm found porphyrin metabolism, purine metabolism, primary bile acid biosynthesis, TCA cycle, alanine aspartate and glutamate metabolism, and the metabolism of xenobiotics by cytochrome p450. Heatmap analyses showed distinct differences in median intensities between the sham-injured and both injured and contralateral-to-injured groups (**Figure 5**).

**Figure 4.**
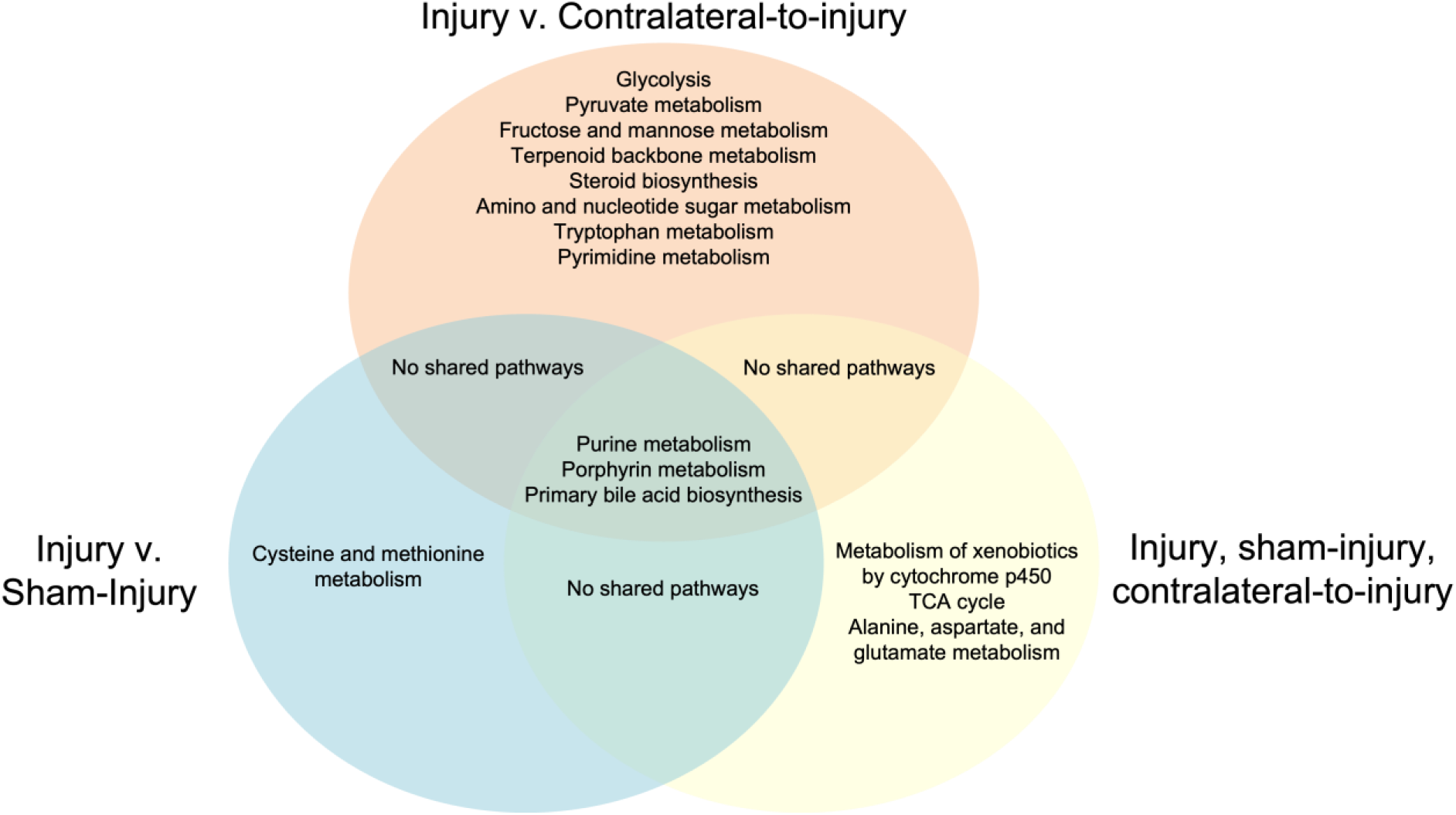
Venn diagram of key synovial fluid metabolic pathways between injured, sham-injured, and contralateral-to-injured limbs.

**Figure 5.**
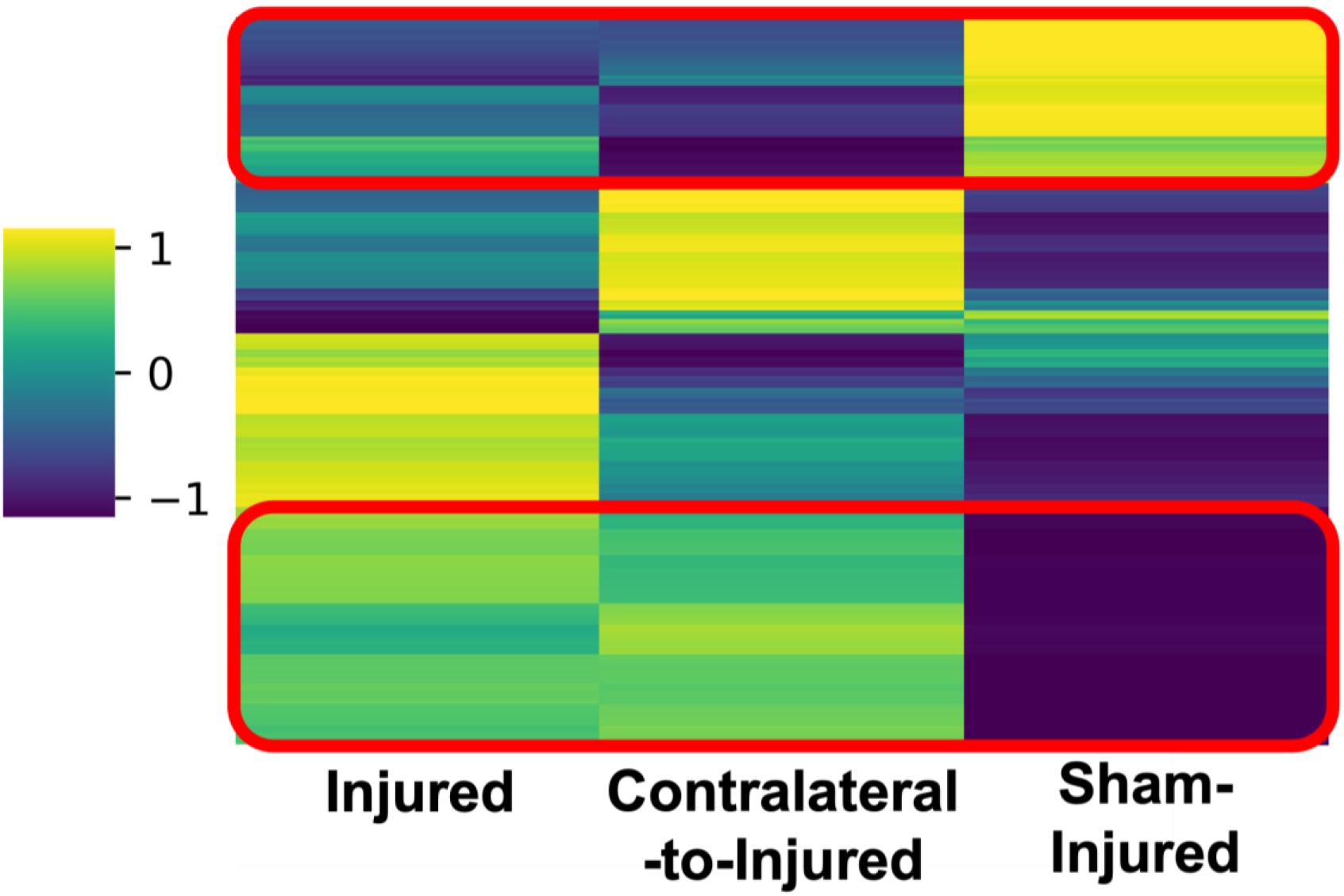
Heatmap analysis of a cluster identified by ensemble clustering of injured, contralateral-to-injured, and sham-injured synovial fluid. Two regions of the heatmap highlighted showing distinct differences between metabolite intensities of injured and contralateral-to-injured, and sham-injured synovial fluid.

#### 3.4.2 Injured and sham-injured synovial fluid have distinct metabolic profiles

Because significant features were detected from ensemble clustering analyses and heatmap analysis confirmed differences in median metabolite intensities between injured, sham-injured, and contralateral-to-injured groups, we compared specific metabolites within these groups. Univariate analysis of injured v. sham-injured synovial fluid revealed 148 significant metabolites. Injured limbs had 98 upregulated and 50 downregulated metabolites compared with sham-injured limbs, after correction for false discovery rate^23^. PCA revealed distinct grouping of injured and sham-injured metabolic profiles apart from one outlier and was associated with 57.9% of the variability. The outlier was an injured limb which clustered with sham-injured limbs in PCA. PLS-DA revealed distinct groupings of injured and sham-injured metabolic profiles and was associated with 31.9% of the variability. Similarly, hierarchical clustering revealed clear grouping of injured versus sham-injured limbs, except for the same outlier detected by PCA analyses.

Clustering validation indicated that 3 clusters was appropriate (**Figure 1D**). The top ten ensemble clustering (**Supplementary Figure 2**) outputs combined with the mummichog algorithm found purine metabolism, porphyrin metabolism, cysteine and methionine metabolism, and primary bile acid biosynthesis.

#### 3.4.3 Injured and contralateral-to-injured limbs have minor differences in synovial fluid metabolic profiles

Univariate analysis of injured v. contralateral-to-injured found no significant metabolic features. Further, PCA analysis indicated similar metabolic profiles for the first 5 components. However, PLS-DA analysis revealed distinct clustering of injured and contralateral-to-injured limbs, with all but one metabolic profile grouping appropriately. Hierarchical clustering of these data revealed no distinct groupings.

Two clusters were deemed optimal from clustering validation. Ensemble clustering (**Supplementary Figure 3**) found purine metabolism, porphyrin metabolism, glycolysis, pyruvate metabolism, fructose and mannose metabolism, amino sugar and nucleotide sugar metabolism, primary bile acid biosynthesis, glyoxylate and dicarboxylate metabolism, terpenoid backbone metabolism, steroid hormone biosynthesis, tryptophan metabolism, pyrimidine metabolism.

## 4.0 Discussion

Early therapeutic interventions may be necessary to slow the progression of PTOA. However, the initial events in the progression toward PTOA are not well understood. In this study, we address several critical gaps of the effect of joint injury on (1) the acute (i.e., 7 days after injury) response of subchondral bone modulus to joint injury, (2) systemic changes (i.e., injured versus contralateral-to-injured limbs) to bone microarchitecture and material properties, and (3) shifts in the synovial fluid metabolome unique to the injured limb.

We found that epiphyseal and subchondral bone microarchitecture undergo bone loss 7 days after joint injury. Previous studies^6–10^ with this injury model in mice also show reductions in epiphyseal bone microarchitecture within 7 to 10 days of injury (**Table 1**). Our study agrees with previous works that epiphyseal bone loss rapidly occurs after joint injury. However, our study finds a 10.8% decrease in BV/TV and a 5.5% decrease in Tb.Th compared to 18-40%, and 18-24% decreases in these measures, respectively^6,8–10^. A potential explanation for these differences is the age of mice used in each study. We chose to study skeletally mature (19-weeks at time of injury) mice, whereas the previous studies utilized skeletally immature mice (10-11 weeks at time of injury). These results demonstrate that early changes to bone microarchitecture may be less profound in the context of skeletal maturity.

**Table 1:**
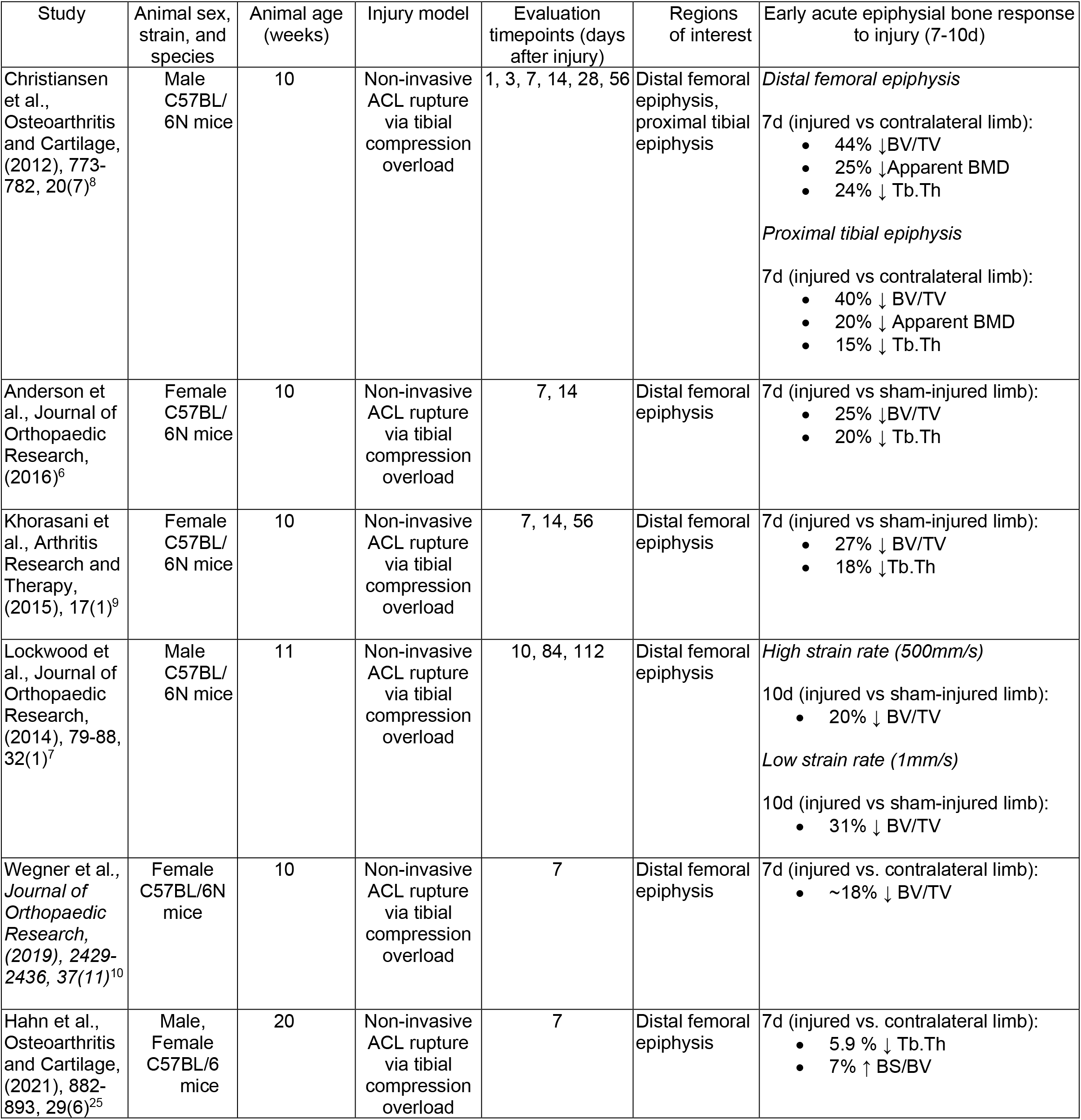
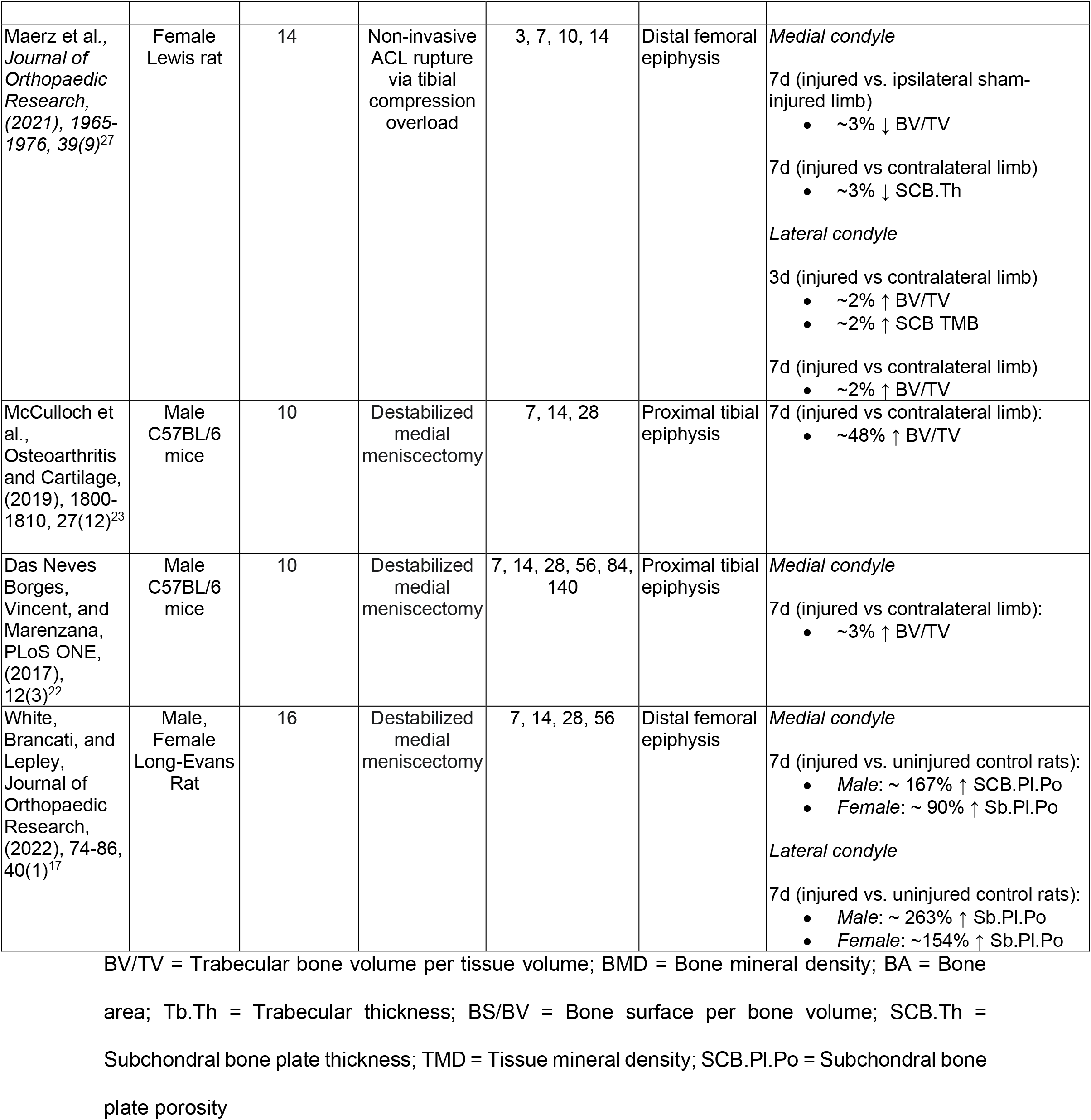
Literature review of early bone response to ACL injury in rodent models

**Table 2:**
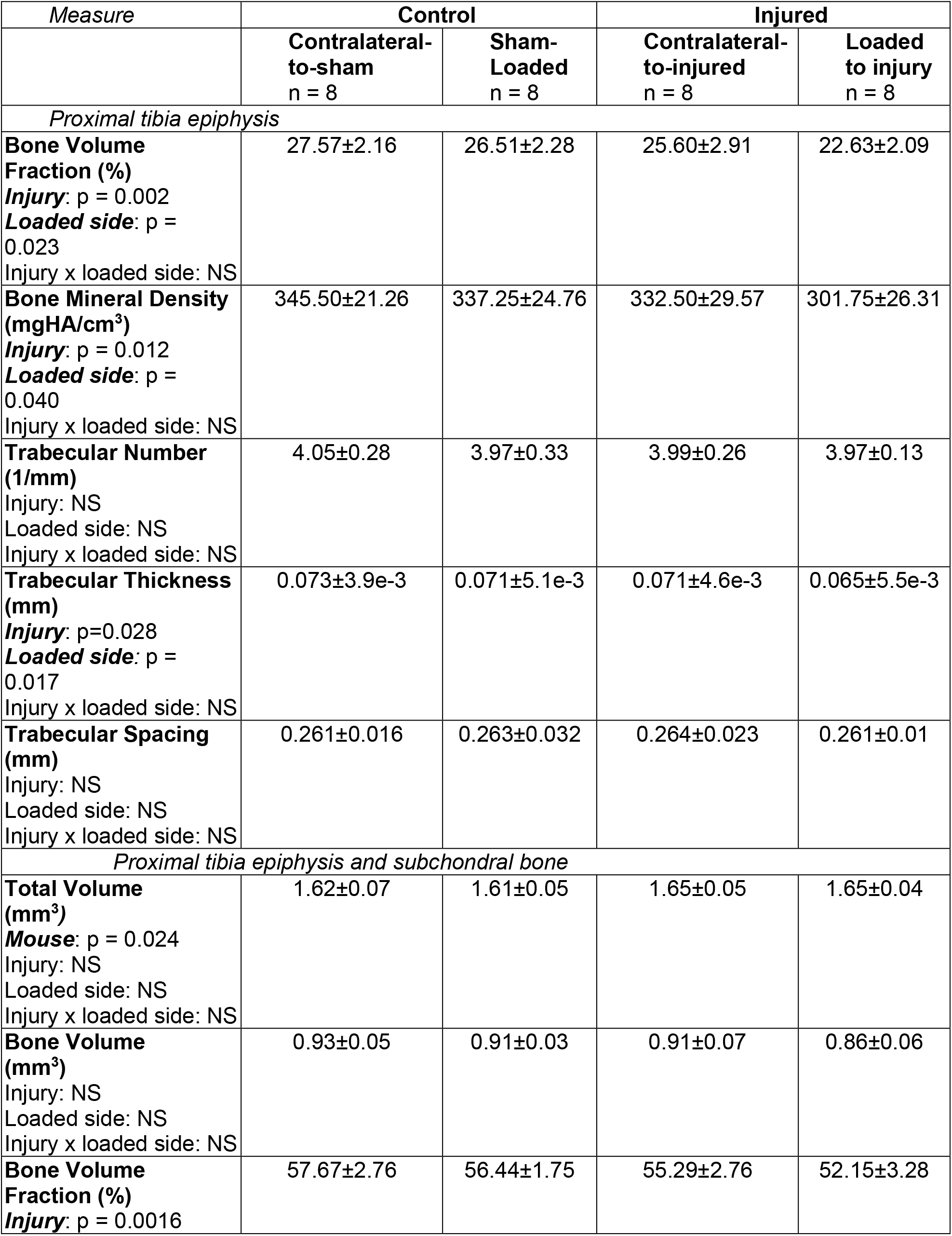
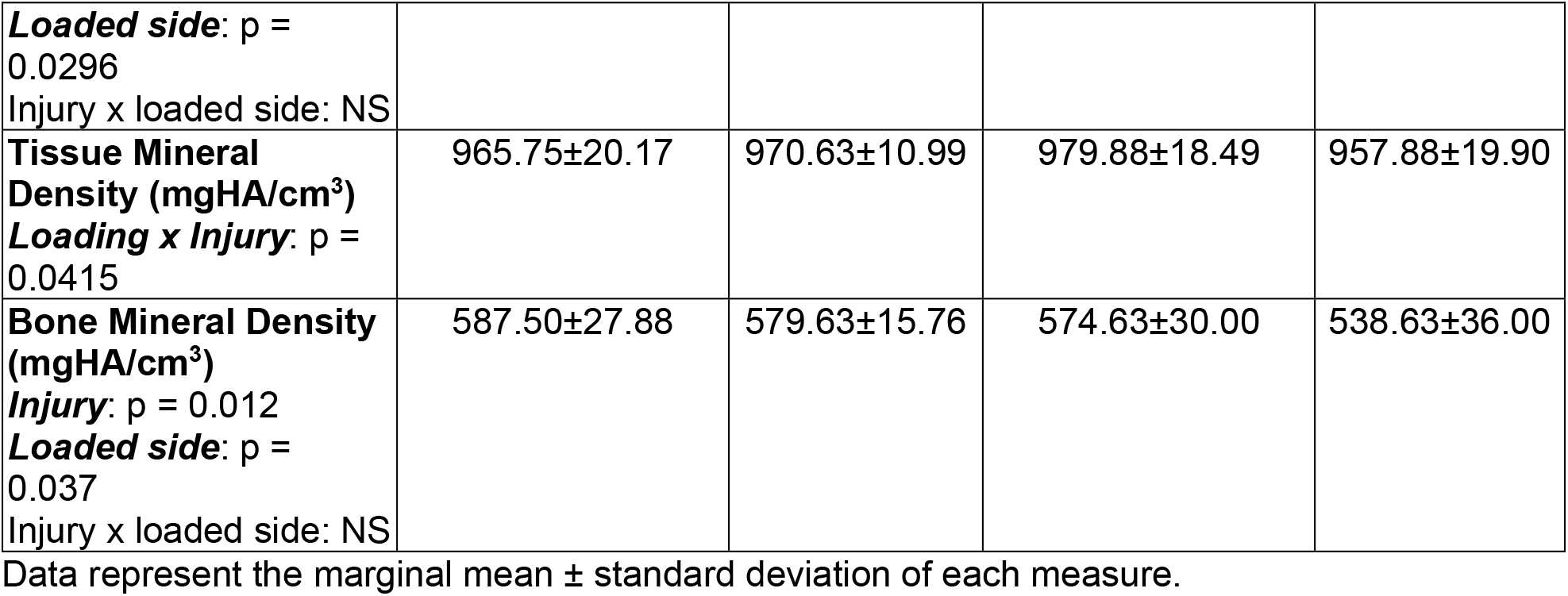
Subchondral bone microarchitecture.

**Table 3:** Subchondral bone modulus, bone formation, and osteoclast number density.

| *Measure* | **Control** |  | **Injured** |  |
| --- | --- | --- | --- | --- |
|  | **Contralateral-to-sham** | **Sham-Loaded** | **Contralateral-to-injured** | **Loaded to injury** |
| *Proximal tibia epiphysis* |  |  |  |  |
| **Indentation Modulus (GPa)**<br>Injury: $p = 0.052$<br>Loaded side: NS<br>Injury x loaded side: NS | 18.33±1.36 | 16.90±3.05 | 19.47±1.41 | 19.39±2.76 |
| **Calcein to Bone Surface (%)**<br>**Mouse:** $p = 0.013$<br>Injury: NS<br>**Loaded side:** $p = 0.005$<br>Injury x loaded side: NS<br>Injury x location: NS<br>Location x loaded side: NS | 46.10±16.35 | 34.54±17.01 | 40.85±13.00 | 36.50±9.08 |
| **Alizarin to Bone Surface (%)**<br>**Limb(Mouse):** $p = 0.001$<br>Injury: NS<br>Loaded side: NS<br>Injury x loaded side: NS<br>Injury x location: NS<br>Location x loaded side: NS | 42.28±18.84 | 33.44±19.82 | 37.63±12.66 | 46.30±23.26 |
| *Distal femur* |  |  |  |  |
| **Osteoclast number per bone marrow area (1/mm<sup>2</sup>)**<br>Injury: $p = 0.017$<br>Loaded side: NS<br>Injury x loaded side: NS | 2.08e-4±2.62e-4 | 1.84e-4±1.27e-4 | 5.53e-4±4.27e-4 | 5.28e-4±3.05e-4 |
| **Osteoclast number per bone surface (1/mm)**<br>Injury: $p = 0.016$<br>Loaded side: NS<br>Injury x loaded side: NS | 5.32e-3±5.64e-3 | 4.26e-3±2.75e-3 | 1.37e-2±1.06e-2 | 1.45e-2±7.77e-3 |

Data represent the marginal mean ± standard deviation of each measure. Mouse numbers per group vary per measure: indentation modulus (injured n = 8, control n = 8); calcein-to-bone-surface and alizarin-to-bone-surface (injured n = 7, control n = 8); osteoclast measurements (injured n = 8, control n = 7).

While it is well-known that bone microarchitecture undergoes a transient decline before the onset of sclerosis at late-stage OA^6–10^, changes to subchondral bone material properties soon after injury remain unknown. This information is needed to contextualize the disruption to the bone loading environment soon after injury. In this study we find that there is insufficient evidence to conclude that subchondral bone modulus changes 7 days after injury. However, we note that our study may have been underpowered with respect to detecting changes in subchondral bone modulus with injury (p=0.052). Understanding how both microarchitecture and modulus change following joint injuries to evaluate the impact of the early response of subchondral bone to joint injury on osteocyte mechanotransduction^40,41^.

Changes to bone structure and material properties can result from altered bone deposition, resorption, or both. Therefore, we assessed osteoclast number density and quantified the epiphyseal mineralizing surface. Our results show an increase in osteoclast number density but not bone formation surface following joint injury. These results demonstrate that the early bone loss in our mouse model is attributable to increased bone resorption alone.

Our experimental design allowed us to ask whether early bone changes following traumatic injuries affected the contralateral limb. Early subchondral bone changes to the injured limb following joint injuries in rodents are well established^6–10,17,22,23,25,27^. However, while the acute inflammatory response after injury has been shown to affect tissues far from the injury site^16^, whether bone changes to the contralateral to injured limb occur remains unknown. We found that bone microarchitecture decreases, and osteoclast number increases for both the injured and the contralateral-to-injured limb. In some cases, the magnitude of the effects of injury and loading on bone microarchitecture measures are similar. Specifically, injury and loaded side have similarly additive effects on early bone loss for epiphyseal BV/TV, BMD, and Tb.Th. These results improve our understanding of mouse models of PTOA because there are clear impacts to the biological and mechanical environments of both knees soon after the joint injury. Future studies should utilize sham-injured mice as controls for each time point to avoid confounding effects from systemic response to injury. Importantly, it is not understood if these changes to the contralateral-to-injured limb are transient or if they later progress towards PTOA as in the injured limb. It is also not known whether other models of PTOA (e.g., DMM, ACLT) produce similar contralateral limb effects at early timepoints, since the surgical incision confounds the early response.

Synovial fluid is integral to joint health, serving as a lubricant for joint articulation and a nutrient source for joint tissues^42,43^. Synovial fluid metabolic analysis revealed distinct differences between injured and sham-injured knees, suggesting synovial fluid metabolism is sensitive to acute joint injury. Metabolomic profiles were also similar between injured and contralateral-to-injured limbs. Our study contributes to a growing body of work that demonstrates that distinct shifts occur in synovial fluid metabolism in the first week post joint injury^25^ and are consistent with shifts in whole joint metabolism^44^. Synovial fluid metabolic changes are driven by dysregulation of cartilage and bone metabolism following joint injuries^19^. Thus, early shifts in synovial fluid metabolism may be key indicators of the first stages of progression to PTOA. Moreover, these changes to synovial fluid metabolism have the potential to contribute to further dysregulation of cartilage and bone metabolism. These findings motivate the need to further study the relationships between synovial fluid metabolism and the progression of cartilage and bone changes after joint injury.

Ensemble clustering analysis of injured v. sham-injured, injured v. contralateral-to-injury, and injured, sham-injured, contralateral-to-injury synovial fluid samples reveals many metabolic pathways of interest (**Figure 4**). Of specific interest for PTOA was cysteine and methionine metabolism which was only dysregulated for injured compared with sham-injured mice (**Figure 4**). A previous study^45^ linked homocysteine (an intermediary of cysteine metabolism) to increased osteoclast activity in-vitro. As reviewed by Delaissé *et al*,^46^ cathepsin K, a cysteine proteinase produced by osteoclasts, is a key proteinase in bone matrix solubilization^47,48^. Furthermore, cathepsin K may also play a role in cartilage degradation^49^. Therefore, our finding of a shift in cysteine and methionine metabolism in the injured joint synovial fluid may be a key metabolic marker for the early catabolic response of the joint after ACL injury.

Our study had several limitations. The non-invasive mouse injury model does not allow us to differentiate between the effects of injurious-level loading and ACL rupture on subchondral bone changes. However, we attempted to decouple these two effects by initially including peak load as a covariate in mixed model analyses of bone responses. There was not a relationship between peak load and any bone measures, so the covariate was removed for subsequent analyses. We were unable to directly confirm ACL rupture for the injured limbs because of the synovial fluid extraction procedure which can inadvertently transect the ACL. However, we verified ACL rupture by the characteristic release in compressive load (**Figure 1F**)^7,8^ and laxity changes compared to the contralateral joint. Previous studies show that the release in compressive load indicates ACL rupture, as verified by later dissection^7,8^. Another limitation is that embedding bone in PMMA significantly stiffens bone tissue compared with the hydrated condition^33,50^. However, preparing the subchondral bone plate for microscale modulus measurement required stabilizing this small structure with an infiltrating embedding material.

In summary, results from this study demonstrate that bone microarchitecture, osteoclast resorption, and synovial fluid metabolic response are all perturbed 7 days after non-invasive joint injury in mice. A potential biomarker of this initial bone catabolic response may be synovial fluid cystine and methionine metabolism, which was only dysregulated in injured knees. Our results implicate a rapidly changing biological and mechanical environment within both the injured and contralateral-to-injured joints that has the potential for influencing the progression to PTOA.

## 5.0 Acknowledgments

This research was made possible by the Department of Mechanical & Industrial Engineering and the College of Engineering at the Montana State University. We appreciate assistance with microCT analyses from the Center for Advanced Orthopedic Studies µCT Core at Beth Israel Deaconess Medical Center as well as assistance with histological sample preparation from Maria Jerome.

## 6.0 Author Contributions

Experimental design: BDH, RKJ, CMH; Data collection: BDH, CD; Data analysis and interpretation: BDH, CD, CMH; Manuscript drafting: BDH, CMH; Revision of manuscript: all authors; Approval of manuscript: all authors.

