## Supplemental Information for "Subchondral bone and synovial fluid metabolomic profiles are altered in injured and contralateral limbs 7 days after non-invasive joint injury in skeletally-mature C57BL/6 mice"

**Supplemental Material**

**Cluster validation and Ensemble Clustering**

Cluster validation uses a minimum spanning tree classification combined with a validation index (**Supplementary** **Equation 1**) to make a data-driven decision on the optimum number of clusters for a given dataset.

$$\boldsymbol{validation index=}\frac{\boldsymbol{cluster compactness}}{\boldsymbol{cluster separation}}$$

(S1)

The optimum number of clusters for a given dataset is the minimum of the validation index. Then, this number of clusters is used to evaluate the hierarchical cluster outputs. Ensemble clustering evaluates multiple hierarchical clustering outputs to determine the percentage of times each metabolite (from 1 to N) clusters with every other metabolite within the data set. Thirteen different hierarchical clusterings were selected for this study because it allows permutation across the single, average, and complete linkage functions and the Euclidean, squared-Euclidean, standardized-Euclidean, and Chebyshev distance measures, along with the Ward linkage function with the Euclidean distance measure (**Figure 3**).

**Ensemble Clustergrams**
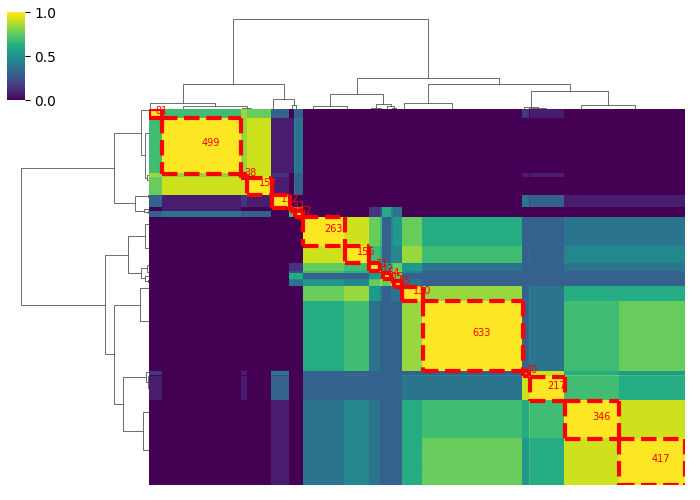

**Supplementary Figure 1** Ensemble clustergram for Injured, sham-injured, contralateral-to-injured comparison

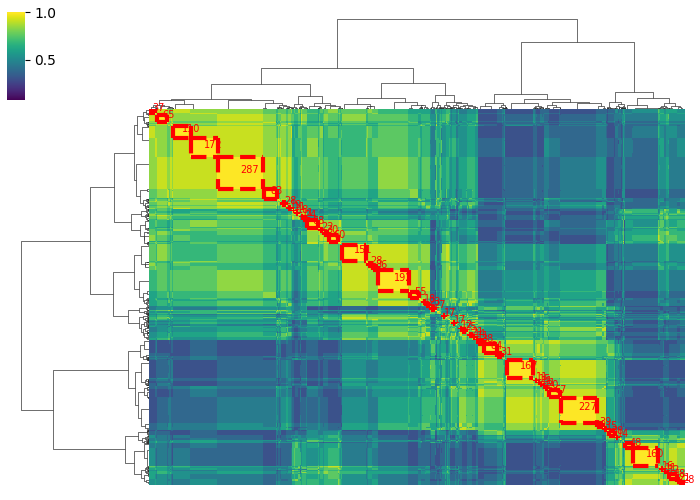

**Supplementary Figure 2** Ensemble clustergram for Injured v. sham-injured comparison

**Supplementary Figure 3** Ensemble clustergram for Injured v. contralateral-to-injured comparison
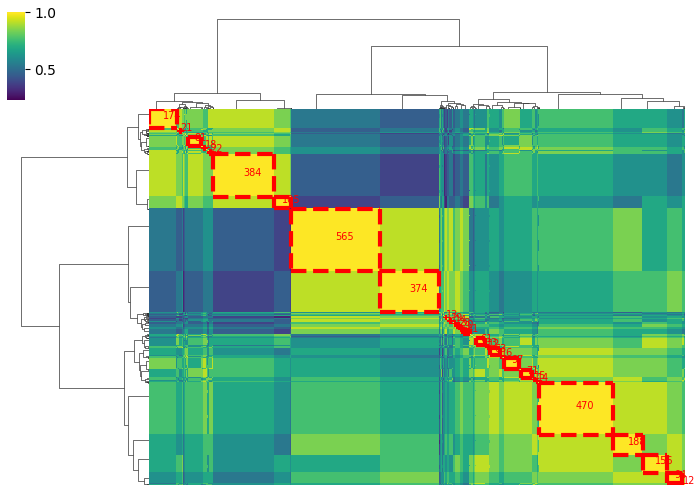

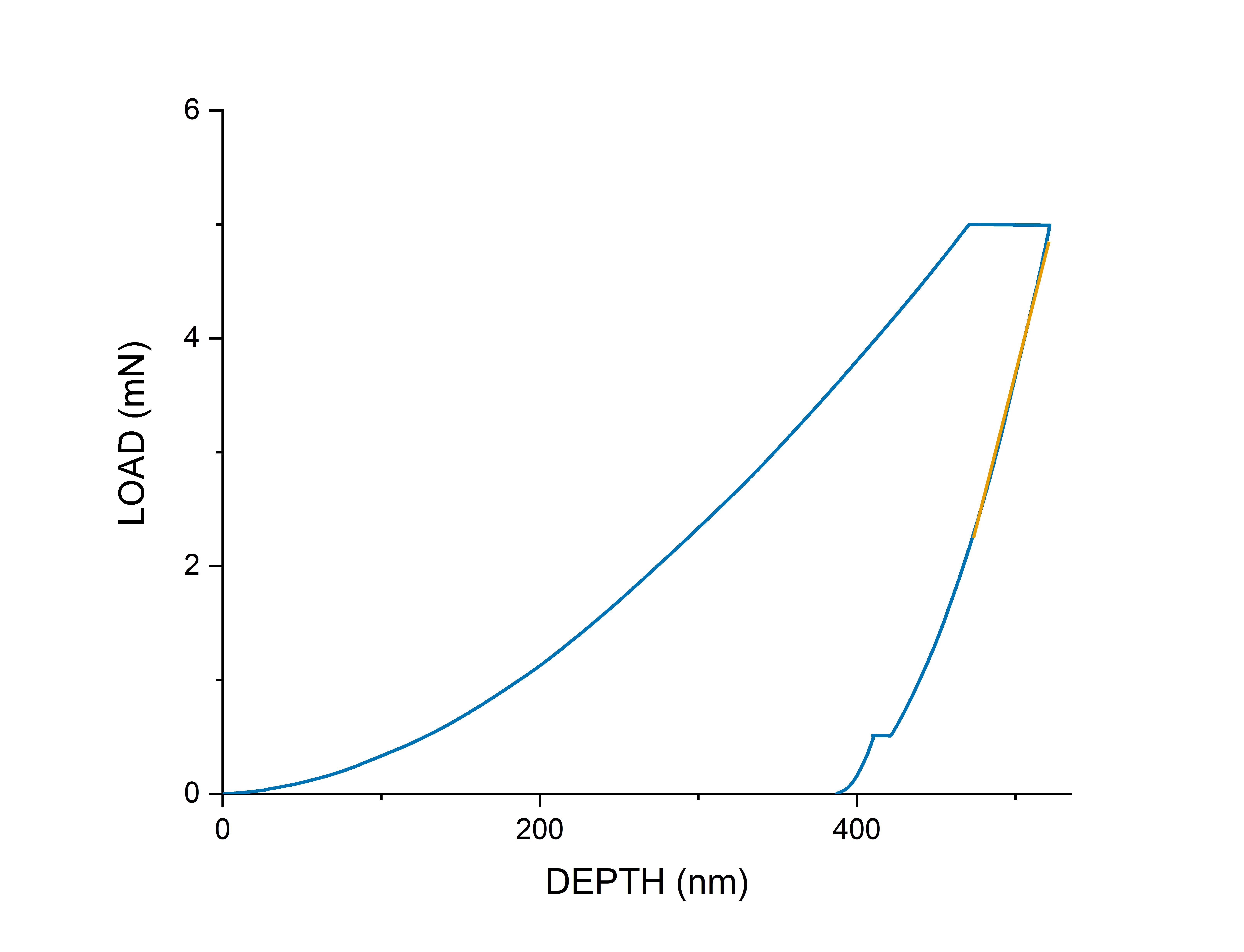

### Supplemental Figure 4 Example nanoindentation curve
